# The Human Dynamic Clamp reveals the fronto-parietal network linking real-time social coordination and cognition

**DOI:** 10.1101/651232

**Authors:** G. Dumas, Q. Moreau, E. Tognoli, J.A.S. Kelso

**Affiliations:** Human Genetics and Cognitive Functions, Institut Pasteur, UMR3571 CNRS, Université de Paris, Paris, (75015) France; Human Brain and Behavior Laboratory, Center for Complex Systems and Brain Sciences, Florida Atlantic University, Boca Raton, FL, USA; Department of Psychology, Sapienza University, Via dei Marsi 78, 00185, Rome, Italy; IRCCS Fondazione Santa Lucia, Via Ardeatina 306, 00100, Rome, Italy; Intelligent Systems Research Centre, Ulster University, Derry, Northern Ireland

**Keywords:** Social cognition, Self-Other Integration, rTPJ, Coordination Dynamics, Virtual Partner Interaction, EEG, Human Dynamic Clamp

## Abstract

How does the brain allow us to interact with others, and above all how does it handle situations when the goals of the interactors overlap (i.e. cooperation) or differ (i.e. competition)? Social neuroscience has already provided some answers to these questions but has tended to treat high-level, cognitive interpretations of social behavior separately from the sensorimotor mechanisms upon which they rely. The goal here is to identify the underlying neural processes and mechanisms linking sensorimotor coordination and intention attribution. We combine the Human Dynamic Clamp (HDC), a novel paradigm for studying realistic social behavior between self and other in well-controlled laboratory conditions, with high resolution electroencephalography (EEG). The collection of humanness and intention attribution reports, kinematics and neural data affords an opportunity to relate brain activity to the behavior of the HDC as well as to what the human is doing. Behavioral results demonstrate that sensorimotor coordination influences judgements of cooperativeness and humanness. Analysis of brain dynamics reveals two distinct networks related to integration of visuo-motor information from self and other. The two networks overlap over the right parietal region, an area known to be important for interpersonal motor interactions. Furthermore, connectivity analysis highlights how the judgement of humanness and cooperation of others modulate the connection between the right parietal hub and prefrontal cortex. These results reveal how distributed neural dynamics integrates information from ‘low-level’ sensorimotor mechanisms and ‘high-level’ social cognition to support the realistic social behaviors that play out in real time during interactive scenarios.

**Significance Statement:** Daily social interactions require us to coordinate with others and to reflect on their potential motives. This study investigates the brain and behavioral dynamics of these two key aspects of social cognition. Combining high-density electroencephalography and the Human Dynamic Clamp (a Virtual Partner endowed with human-based coordination dynamics), we show first, that several features of sensorimotor coordination influence attribution of intention and judgement of humanness; second, that the right parietal lobe is a key integration hub between information related to self- and other-behavior; and third, that the posterior online social hub is functionally coupled to anterior offline brain structures to support mentalizing about others. Our results stress the complementary nature of low-level and high-level mechanisms that underlie social cognition.

## Introduction

Much of our social life consists of interactions with others. Despite their essential role, social interactions still remain the “dark matter” of social neuroscience: the field is in urgent need of studies that embrace the reciprocal and real-time nature of social coordination (Hari and Kujala, 2009; Dumas, 2011; Hasson et al., 2012; Konvalinka and Roepstorff, 2012; Schilbach et al., 2013; Hari et al., 2015). Overcoming the methodological challenge of bringing a true interactive context into the laboratory, some innovations have allowed behavioral tracking, brain recording and stimulation of multiple participants in interaction (Montague et al., 2002; Babiloni et al., 2006; Tognoli et al., 2007; Dumas et al., 2010; Funane et al., 2011; Moreau et al., 2016; Chen et al., 2017; Dikker et al., 2017; Hirsch et al., 2017; Novembre et al., 2017; Era et al., 2018b). Such investigations have exposed the neural underpinnings of our propensity to interact with others during sensorimotor coordination (Tognoli and Kelso, 2015).

Social neuroscience studies that investigate the interaction between multiple participants face the challenge of having more sources of unconstrained variance than experimental paradigms with single participants do. To overcome this limitation while at the same time maintaining the essential reciprocal and dynamical nature of social interaction, we use the Human Dynamic Clamp (HDC) or Virtual Partner Interaction (VPI) paradigm which enables direct experimental control of one of the social partners, as well as the coupling between them. The HDC consists of a human interacting reciprocally with a virtual partner (VP), the design of which is based on an empirically grounded computational model of human coordination dynamics (Kelso et al., 2009; Dumas et al., 2014a; Kostrubiec et al., 2015). Both the intrinsic dynamics of the VP and its coupling to the human can be manipulated in real-time thereby enabling parametric exploration of the relationship between humans and interaction-capable surrogates. Analogous to its cellular counterpart (Prinz et al., 2004) the HDC or VPI paradigm offers a computationally precise way to approach the complexity of real-life social interaction while at the same time maximizing experimental control.

Divergent theories of social cognition have risen over the years. On the one hand, cognitive theories have focused on ‘top-down’ processes (Sebanz et al., 2006), mentalizing and theory of mind (Frith and Frith, 2012); on the other hand, sensorimotor theories have focused on the spontaneously self-organizing nature of coordination (Oullier et al., 2008; Coey et al., 2012), mirroring (Rizzolatti and Sinigaglia, 2016), and sensorimotor coupling (Hari and Kujala, 2009). The fully bidirectional nature of social behavior in terms of the perception-action has stressed the reciprocal relationship between levels of analysis and direction of information flow (Kelso et al., 2013). Previous behavioral studies have shown that whether humans perceive VP behavior as cooperative or competitive is modulated by coupling strength (Drever et al., 2011). Interestingly, however, automatic imitation processes (i.e. visuo-motor interference) seem to be present in both competitive and cooperative scenarios (Era et al., 2018a). Using the Human Dynamic Clamp, successful sensorimotor coupling (i.e. stably coordinated movements) and attribution of humanness to the virtual partner have been associated with increased emotional arousal (Zhang et al., 2016). Moreover, under certain conditions, such as competing task requirements, humans spontaneously endow the VP with intention and goal-directedness (Kelso et al., 2009; Drever et al., 2011).

The present work aims to elucidate so called top-down and bottom-up perspectives of social behavior using real-time interaction between a human and a Virtual Partner in conjunction with neurodynamical analyses of spatially-resolved high-density EEG recordings. A benefit of using the HDC is that the evolving brain dynamics can be correlated with both human and VP movements as well as their coordination. In addition, the paradigm enables an evaluation of how humans reflect on their interaction with the VP and assess their (virtual) partner’s humanness and intention. The goal is to expose the brain circuitry hypothesized to relate real-time coordination and ongoing cognitive and emotional aspects of social behavior.

## Material and Methods

### Participants

20 volunteers, 12 males and 8 females, aged between 18 and 33 years (Mean=24.2, STD=4.5) took part in the study. All were right-handed (Edinburgh Handedness Inventory) with reported normal or corrected-to-normal visual acuity and without self-reported history of neuropsychiatric disease or movement disorder. Participants provided informed consent prior to the research. The study was approved by the Internal Review Board at Florida Atlantic University and conformed to the principles expressed in the Declaration of Helsinki.

### Material and apparatus

Participants were seated in a dark Faraday room, with the ulnar side of the right forearm resting against a U-shaped support (21.5 × 8 × 4 cm) positioned parallel to a table (Fig. 1A). Participants supported their right hand by grasping a vertical wooden cylinder (4.5 × 3 cm), leaving only the right index finger in extension. The hand was oriented in the sagittal plane and the distal part of the index finger was inserted into the circular orifice (2 cm diameter) of another wooden block. The latter was connected through two metallic bars to a vertical, freely rotating metallic stem (18 cm length) whose angular displacement was captured by a linear potentiometer placed atop. The entire arrangement constituted a manipulandum, which was fixed on top of a Plexiglas box (30.5 × 31.5 × 20 cm), positioned to the right of a screen, about 50 cm away from the midline of the participant. The manipulandum restricted the movement of the index finger to the horizontal plane and allowed a full-range of friction-free flexion-extension motion about the metacarpophalangeal joint.

**Figure 1.**
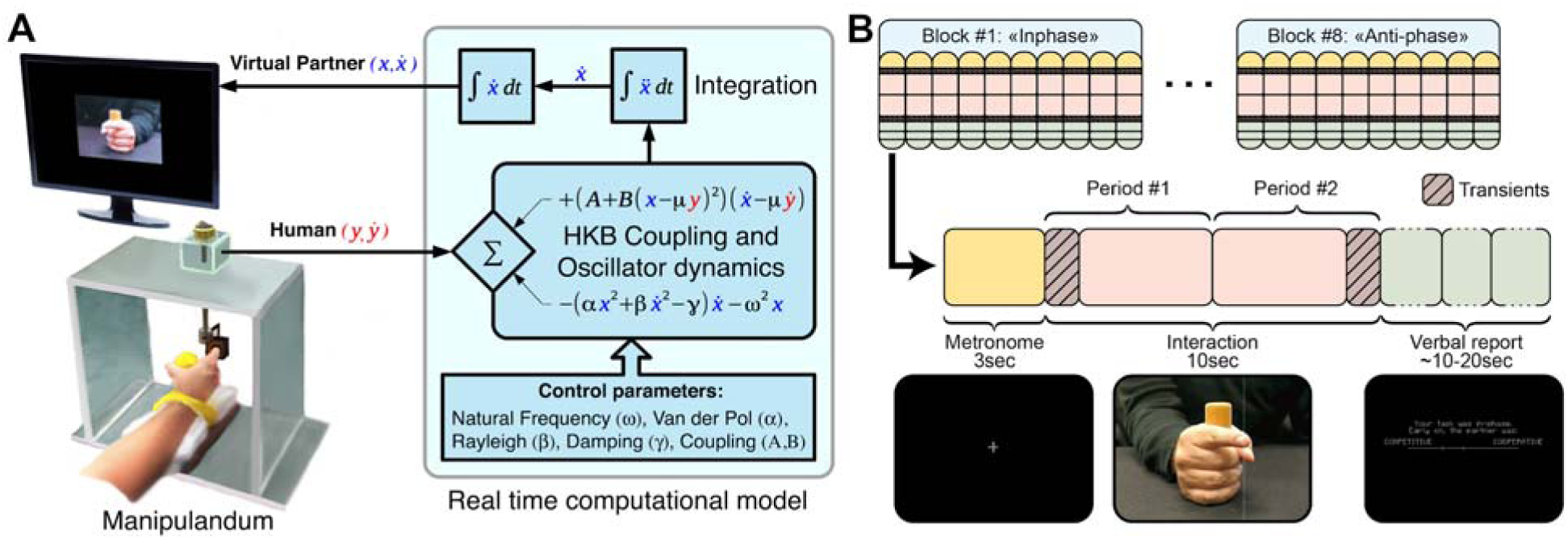
Schematic of the Human Dynamic Clamp paradigm. (**A**) Schematic of the HDC experimental apparatus. Human participants coordinate finger movement with a virtual partner (VP) displayed on a screen. Movements are digitized and fed into a computer, where the HDC software computes in real time the corresponding position of the VP following the Haken-Kelso-Bunz (1985) model. The key parameters of the HKB model are the VP’s intrinsic frequency (ω=1.6Hz), and terms that control movement shape and various dynamical properties of the oscillator for behavioral realism (Van der Pol α=0.641, Rayleigh β=0.007095, and damping γ=12.457), the coupling (A = −0.5 and B = −0.25), and the in-phase/anti-phase preference (μ=+/-1). (**B**) Summary of the experimental paradigm (a non-verbal Turing test) showing the structure of trial blocks (see text for details).

The output of the potentiometer was sampled at 1000 Hz in step with the EEG, using a National Instruments A/D converter. The signal was down-sampled offline to 500 Hz for online computational efficiency, and used as a continuous input into a computer, which ran the Human Dynamic Clamp program implemented on C++ using the cross-platform IDE Code Blocks and the open source library OpenFrameworks (code available at https://github.com/GHFC/SoNeTAA). A java script version of the Human Dynamic Clamp is available on GitHub: https://github.com/crowd-coordination/web-vpi and can be tested online at http://www.morphomemetic.org/vpi/. The velocity of the human finger was numerically computed using a 3-point differentiation algorithm and, together with position data plugged into the VP equation (see Dumas et al., 2014 for details). The differential equation returned instantaneous VP acceleration, which was integrated using a 4th order Runge-Kutta method at 500 Hz to provide VP velocity and position. We compared digital timestamps and analog triggers with an oscilloscope to ensure that a maximum delay of 2 ms occurred between data acquisition and computation of the model output (0.3% of typical movement cycle length).

To create the animation of VP finger movement used in the experiment, a series of position-indexed images was created and stored (see Kelso et al., 2009). The images were recorded with a high-speed camera while a human male hand produced flexion-extension finger motions in the horizontal plane. A complete cycle of movement provided 119 images (17 × 13 cm) indexed by their position. The instantaneous position of the VP, as computed in real time during the experiment, was iteratively used to select one of the 119 position-indexed images, which was displayed in the center of the screen (59 cm diagonal). The screen animation was refreshed at 100Hz during the experiment and looked just like an ordinary video display of a real finger in repetitive motion. An auditory tone of 440 Hz and 0.1 s duration was used as a pacing signal to constrain the initial frequency difference between human and VP, and to minimize inter-trial and inter-subject variability accordingly.

### Procedure

Participants interacted with the VP during a session composed of 80 pseudo-randomized trials (Fig 1B). During the instruction phase, the experimenter demonstrated the ability of a second manipulandum, placed in participant’s view just outside the EEG chamber, to provide real-time control of finger movement. The addition of this manipulandum not only assisted in conveying task instructions but also provided a basis for the participant to attribute observed finger movements to another human. Trials were arranged in blocks of 10 trials, during which participants were tasked to coordinate their movement either in-phase or anti-phase with the VP. The order of blocks was counterbalanced across participants.

Each trial was composed of three periods (Fig. 1B): Pacing (3s), Interaction (10s) and Self-report. Participants were instructed to maintain coordination with the VP during the Interaction according to block-wise instructions (i.e. in-phase or anti-phase). Before each trial, a brief screen presentation indicated the instruction of the current block followed by a fixation-cross at the center of the screen. Then, the Pacing period started, and an auditory tone cued the required movement frequency (1.6Hz). Participants were instructed to produce peak flexion on each beat while fixating upon the cross, and then to maintain the frequency throughout the rest of the trial. As soon as the pacing signal was turned off, the Interaction phase began, and the moving VP finger appeared on the screen. The VP also had an intrinsic frequency of 1.6Hz and a phase randomized and locked relative to the auditory metronome. The VP was randomly assigned a cooperative or competitive behavior for two halves of each trial (i.e. 4s sub-period), giving 4 pseudo-randomized types of trials: cooperation throughout, competition throughout, switch from cooperation to competition, and finally switch from competition to cooperation. After the Interaction period, rating scales were presented on the screen. Participants reported the degree to which the VP was cooperative or competitive early and later on during the trial, and finally judged the humanness of the VP (i.e. if they thought the partner was a human or a machine). All verbal reports were done using a scale on the screen which was controllable by the finger motion of participants. After each choice was dialed on the manipulandum, participants employed a short acquiescing sound to signal validation of their selection without talking, and the experimenter moved the program to the next trial. The nasopharyngeal sound (resembling /_L_/ but without the labial component) had previously been established to minimize muscular and movement artifacts at the sites of EEG recording.

### Behavior analysis

Potentiometer signals corresponding to human finger displacement and position of the VP’s finger were mean-centered, detrended, low-pass filtered using a second order dual-pass Butterworth with a cutoff frequency of 20 Hz and normalized. After this preprocessing, we quantified the coordination between human and VP by calculating the continuous relative phase (RP) between their movements (phase estimated with continuous Hilbert transform). To avoid transients and thus separate the inhomogeneous neural activity occurring at onset and offset of the interaction, we removed the first and last seconds of interaction, leaving 8 s of each trial for analysis. For each of the two halves of the trials, mean RP and the corresponding circular variability were calculated, in order to assess the produced pattern (e.g. in-phase, anti-phase or some other pattern) and its stability, respectively. For each participant, verbal reports of humanness and cooperativeness were Z-score normalized, and performances were quantified through two indices: a phase coordination score equal to the normalized absolute difference between ongoing relative phase and the RP corresponding to the task condition (i.e. 0 rad for in-phase, and pi rad for anti-phase); an intention attribution score equal to the normalized difference between perceived cooperativeness and real VP behavior, thus quantifying if the human participant was able to perceive VP’s helpfulness (or not) toward achieving his/her goals. Behavioral variables were analyzed through a repeated measures 2×2×2×2 ANOVA in JASP (JASP Team, 2018) having as factors Trial-Part (First-Half/Second-Half), Behavior of the VP (Cooperative/Competitive), Transition during the trial (Yes/No), and Task (in-phase or anti-phase). Results were corrected for multiple comparisons using Bonferroni tests (significance level at *p < 0.05*).

### High-density EEG recording

Various neurophysiological markers have been implicated in social processes, including fronto-central and occipito-temporal Theta (4-7Hz; Dumas et al., 2012; Bramson et al., 2018; Moreau et al., 2018), several types of fronto-parietal Alpha (8-13 Hz; Tognoli et al., 2007; Konvalinka et al., 2010; Novembre et al., 2014; Tognoli and Kelso, 2015), rolandic Beta (Ménoret et al., 2013), and parietal Gamma (>30Hz; Dumas et al., 2010, 2012). Accordingly, EEG analyses were conducted across all frequencies and without predetermined judgment regarding regions of interest. Via this strategy we aimed to: 1) quantify how the dynamics of social coordination with a virtual partner affects human subjective reports (cooperation∼competition); 2) identify neuromarkers (local oscillations, transiently coupled functional networks) associated with sensorimotor coupling (i.e. integration of movements from self and other); and 3) investigate the neural correlates of humanness and intention attributed to a virtual partner.

The experiment was conducted in a sound-proof Faraday chamber. High-density EEG was recorded using 128 channel EEG caps with Ag-AgCl electrodes (Falk Minow Services, Germany) arranged according to an extension of the 10-20 system (Jasper, 1958; Oostenveld and Praamstra, 2001). The signals were fed to an amplifier (Synamp2; Neuroscan, TX) equipped with high-level port to ensure recording of triggers from the HDC software and to obtain temporally precise analysis of brain-behavior dynamics. EEG signals were measured with the respective ground located on the left shoulder and referenced at the Cz electrode. Impedances were maintained below 10 k∧. The signals were analog filtered (Butterworth, band pass from 0.05 Hz (−12 dB per octave) to 200 Hz (−24 dB per octave), amplified (gain of 2,010) and digitized at 1,000 Hz with a 24-bit ADC in the range ±950 µV (vertical resolution of 0.11 nV).

### Skin Potential Response (SPR)

Emotional arousal was quantified with a bipolar montage of two passive Ag/AgCl electrodes capturing sympathetic changes (one on the left palm and one on the left epicondyle as a reference, both placed on the immobile hand). We extracted the SPR normalized magnitude by following the method described in Zhang et al. (2016) (where the main results from this experiment with respect to emotional arousal are reported).

### Artifact correction and data preprocessing

Following visual inspection, any noisy EEG channel was marked as bad (average=4.5, min=0, max=10) and interpolated using a spherical spline algorithm (Perrin et al., 1989) with an interpolation order m=3, a Legendre polynomial order n=50, and a regularization parameter λ = 10e-8 (Kang et al., 2015). Correction of eye blink artifacts in the EEG data was performed using a classical PCA filtering algorithm (Wallstrom et al., 2004) on 800ms windows with 400ms overlap. A Hamming window was used to control for artifacts resulting from data splicing. EEG signals were then visually checked to exclude from analysis all trials contaminated by residual eye blinks, unwanted swallowing, coughing, or movement artifacts. Following correction, EEG data were re-referenced to a common average reference (CAR). More than 70 artifact-free trials were obtained for all participants and there were no differences in overall quality of the data and number of unrejected trials per condition.

### Single-trial EEG source estimation

Source reconstruction was performed with the free open-source application Brainstorm (http://neuroimage.usc.edu/brainstorm; (Tadel et al., 2011). Sensors were registered for each participant using head points and fiducial landmarks (nasion and preauricular points) digitized with a *Polhemus* Isotrak system. and projected on the scalp surface of the standard Montreal Neurological Institute (MNI) template space (Holmes et al., 1998). The lead field was then computed using a boundary element model in OpenMEEG (BEM) (Kybic et al., 2005; Gramfort et al., 2010) with a cortical surface tessellated with 2000 vertices. A noise covariance matrix was estimated from a 2 min resting state condition. The inverse solution was estimated for each individual using a standardized Low-Resolution Brain Electromagnetic Tomography method (*sLORETA*) (Pascual-Marqui, 2002) with unconstrained source orientation. Thus, cortical sources were estimated at each vertex of the cortex surface with three orthogonal dipolar sources.

### Brain analyses

The estimated cortical source dynamics were then processed for each trial (Fig. 1B) by taking the Interaction period without the first and last one sec. transients, since brain activity was expected to be non-stationary near these boundaries. The remaining 8 sec. were tapered with a one sec. Hamming window, and discrete Fourier transforms (DFT) used to estimate spectra at the source level. We used Student T-test with False Discovery Rate (FDR) correction for the statistical analyses of power modulations. Reports of anatomical regions followed the Tzourio-Mazoyer atlas (Tzourio-Mazoyer et al., 2002).

For functional connectivity, following Maris et al. (2007), the coherence between all pairs of sources was calculated for frequency bands of interest (see table S1) according to the formula:

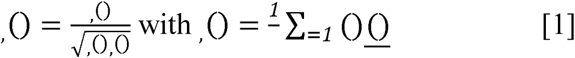

where and are the sources index, () is the cross-spectrum, is the number of windows (i.e. equals to 8), () is the complex Fourier component at frequency for source and__denotes complex conjugate.

**Table 1.**
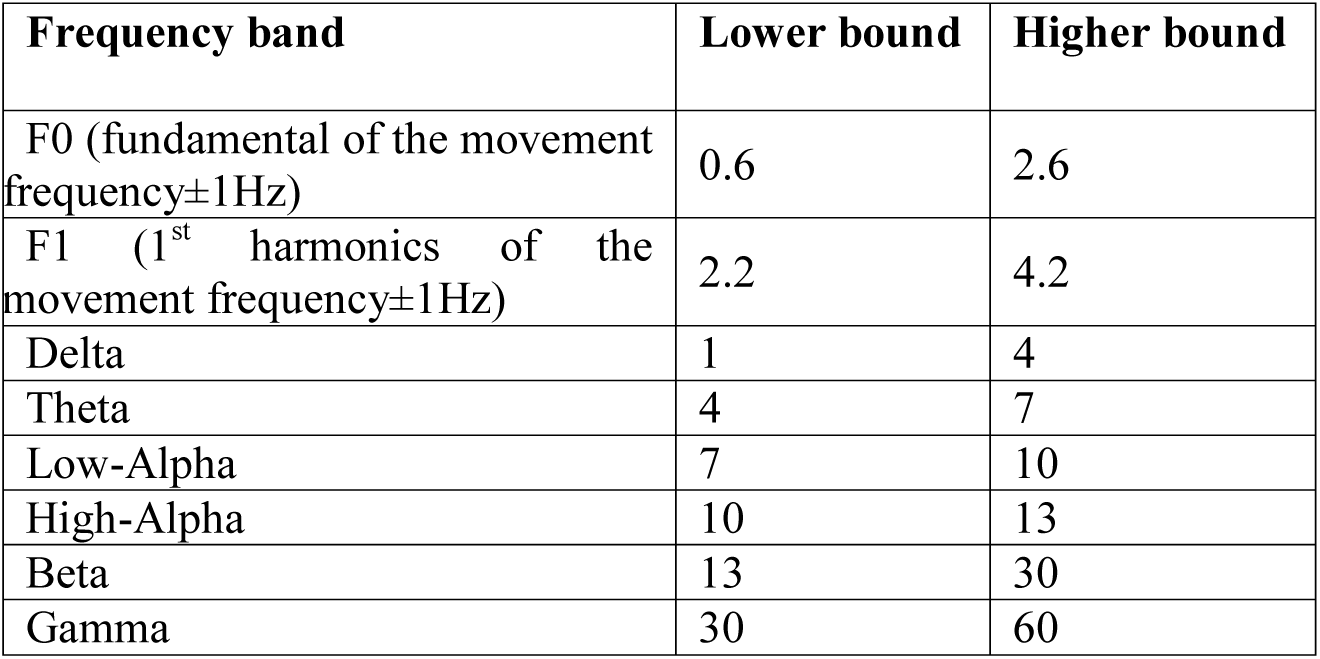
Frequency bands of interest with their respective lower and higher bounds. All values in Hz.

We then averaged the power and coherence values across trials and eliminated potential statistical bias due to the non-Gaussian distribution of coherence values and unequal sample sizes by using Z-Coherence (Maris et al., 2007).

The Z-Coherence is defined as:

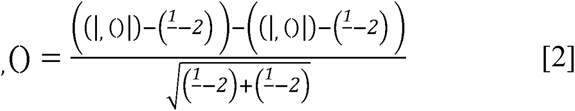

where and are the degrees of freedom in conditions A and B, () and, () are coherences in conditions A and B between sources and at frequency. The sign of Z indicates whether coherence in condition A is higher (positive) or lower (negative) than in condition B. To target large scale brain dynamics links with high-level cognition, we tested whether attribution of humanness and cooperative/competitive actions from the VP were associated with whole-brain network connectivity variations.

### Brain-behavior analyses

We also computed coherence between movement velocity and reconstructed cortical sources. In this case, the formula described for neural coherence in eq.1 still holds except that the source is not another cortical source but either the human or the VP movement velocity. To compute the Z-Coherence, we used as control condition (i.e. condition B, in equation 2) a phase scrambled behavior. We averaged the coherence maps over participants and contrasted the two conditions.

### Statistical tests

Student t tests were computed for both Power and Z-Coherence comparisons. All difference maps were thresholded at *p* < 0.05 by using group-level (n = 20), two-tailed paired permutation tests, which randomly exchanged the estimated values of coupling between conditions for each participant. We used exhaustive permutations (2^20) to estimate the empirical distribution under the null hypothesis of no difference between the two conditions (Pantazis et al., 2005; Maris and Oostenveld, 2007). For cluster-based statistics, the connectivity matrix across the vertices was extracted from the BrainStorm MNI mesh previously used for source reconstruction analysis (i.e. vertices that are part of the same face of the mesh are considered as neighbors). The alpha level was adjusted to the maximum statistical distribution to control for type I family wise error rate (FWER) due to multiple comparisons over the entire brain surface. Unless otherwise stated, all coherence and power plots presented here were obtained with group-level (n = 20) statistical inference based on paired permutation tests. All randomizations were done for rejection of the null hypothesis and to control the false alarm rate at *p* = 0.05.

## Results

### Behavior

Behavioral analysis focused on verbal reports (accuracy in cooperative or competitive intention attribution) and coordination measures (stability of the relative phase during the interaction).

### Participants accurately judged partner’s intention

During the coordination task, both partners had to settle on a common frequency while being instructed to coordinate with each other either in-phase or anti-phase, two modes known to be stable (Haken et al., 1985; Kelso, 1984). In agreement with an experimentally chosen parameter of strong coupling from the VP, the relative phase was mostly stable and followed the intention of VP.

After each trial, participants were asked to judge the intention of the partner, that is, was (s)he cooperative or competitive (see Figure 2). Overall, they correctly attributed VP’s intention 80.3% of the time (*p* < 0.001, permutation test against chance; 10.6% false-cooperation, 9.1% false-competition) and their verbal reports were successfully modulated by the behavior of the VP during the interaction. The 2×2×2 repeated-measures ANOVA (VP Behavior [Cooperative/Competitive] by Transition [Yes/No] by Task [In-phase/Anti-phase]) on subjective reports of cooperativeness revealed a main effect of VP behavior (F(1, 19) = 579.49, *p* <0.001, η^2^ = 0.63) showing that the VP was rated more cooperative in the Cooperative condition compared to the Competitive one. The Transition factor also reached significance (F(1, 19) = 4.25, *p* = 0.04, η^2^ = 0.005), showing that a change in VP behavior during the trial resulted in the VP being rated less cooperative. Finally, there was a main effect of Task (F(1, 19) = 9.71, *p* = 0.002, η^2^ = 0.011), as well as an interaction between VP behavior and Task (F(1, 19) = 10.8, *p* = 0.001, η^2^ = 0.012). Post-hoc tests revealed that VP was judged less cooperative for the Task anti-phase than in-phase (t (19) = −3.4, *p* < 0.001), especially when the trials started with a cooperative VP (Fig. 2).

**Figure 2.**
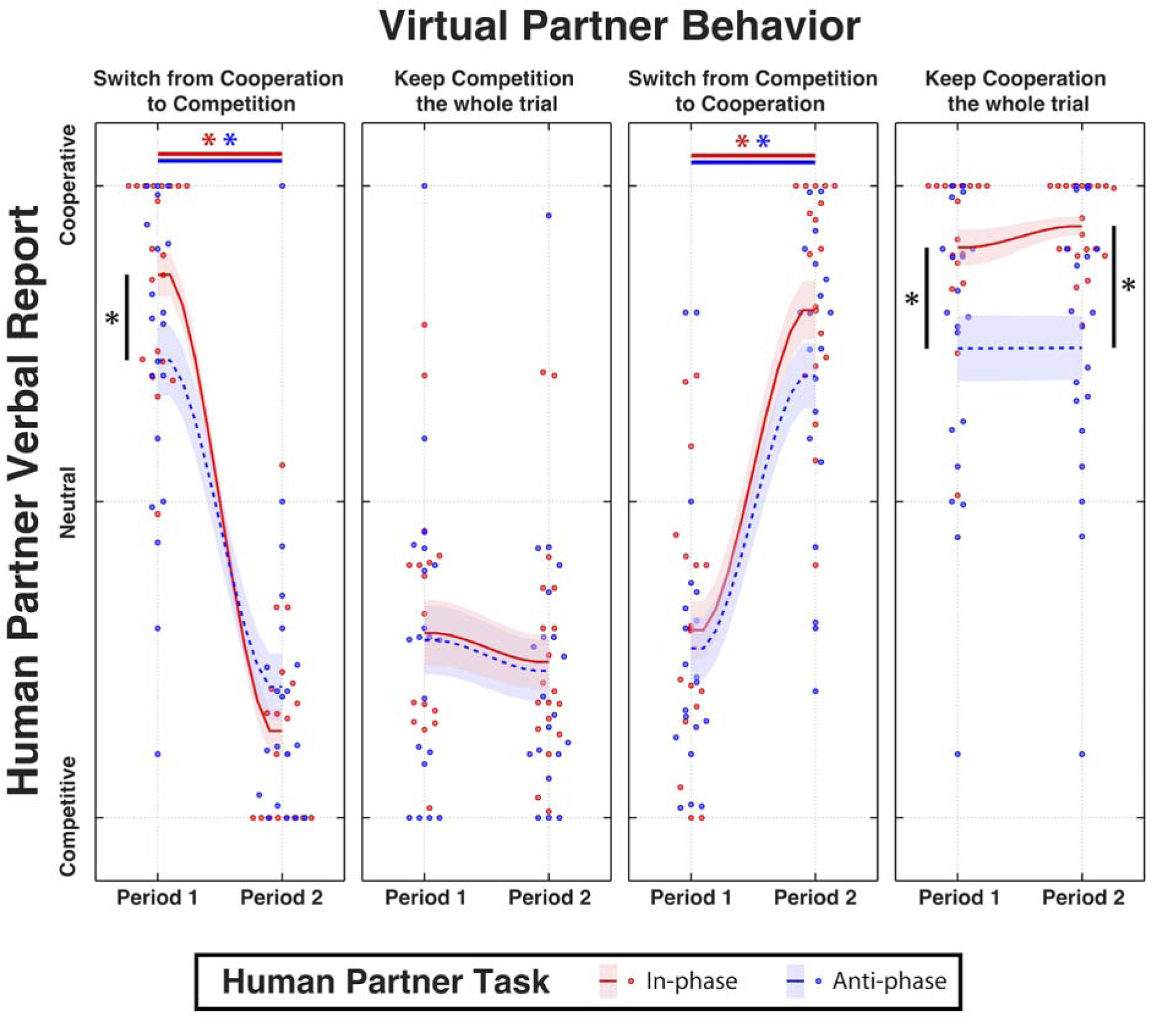
Judgment of cooperation and competition between human and VP. Verbal reports show how participants perceived the VP across the different conditions. Red and blue plots represent respectively the in-phase and anti-phase tasks, shaded areas represent standard errors and stars indicate statistically significant differences (*p <* 0.05, corrected). See details in text. One notable exception occurred when the human task to move in-phase led to a VP deemed less cooperative than when the human was tasked to move anti-phase (Fig. 2, left panel). This happened during the second part of trials with a competitive VP intending anti-phase, following initial segments when VP cooperated in-phase (“treachery”). In such cases, VP was deemed the most competitive of all (Fig. 2, left panel). Overall, the results suggest already that sensorimotor coupling and intention attribution are linked: the better able were participants to coordinate together, the better aware they were of their partner’s intention.

### Stability of VP intention and coordinative stability mediates social cognition

The 2×2×2×2 repeated-measures ANOVA (VP Behavior [Cooperative/Competitive] by Transition [Yes/No] by Task [In-phase/Anti-phase] by Trial-Part [First-Half/Second-Half]) on relative phase stability revealed that coordinative stability depended: a) on the cooperativeness of the Virtual Partner (F(1, 19) = 42.45, *p* < 0.001, η^2=^ 0.104) with competitive being less stable than cooperative behavior; b) on the organization of the trial, with the first half of the trial less stable than the second (F(1, 19) = 15.97, *p* < 0.001, η^2=^ 0.039); and c) on the presence of a transition in VP behavior, with the presence of a transition resulting in less stability (F(1, 19) = 19.34, *p* < 0.001, η^2=^ 0.048). The Transition x Trial-Part interaction (F(1, 19) = 20.47, *p* < 0.001, η^2=^ 0.050) indicated that relative phase stability was lowest in the second half of the trials, in cases where VP switched behavior at mid-trial (*ps <* 0.001). Note that in all cases, stability was assessed in each half of the trial and after a transient to discard effects from the appearance of the partner and a switch in VP intention. Interestingly, accurate intention attribution was correlated with the stability of the interaction (r = 0.51, *p* = 0.02), pointing toward a link between the pattern of sensorimotor coordination and socio-cognitive assessment of the other.

### Cooperative Virtual Partners were judged more human

Consistent with earlier reports that endowed Virtual Partners with intentionality (perceived as the VP trying to trick the human to achieve goals opposite to his/her, Kelso et al., 2009), we found that (objectively) cooperative VPs scored higher in humanness ratings than competitive ones (Wilcoxon rank test Z = 4495.5, *p* < 0.002). Despite this prominent dependency on objective variables of cooperativeness, the humanness rating was independent of subjective judgement of cooperation. Indeed, subjective ratings of cooperativeness in both periods of interaction did not show any differences between trials associated with judgments of humanness (Wilcoxon rank test with *p* = 0.6 for the first part of the trial, and *p* = 0.84 for the last). Similarly, trials with both parts judged as cooperative did not demonstrate differences in humanness rating compared to trials with both parts judged as competitive (Wilcoxon rank test, *p* = 0.13). In contrast to intention attribution, judgment of humanness did not depend on relative phase stability (Wilcoxon rank test, *p* = 0.52). However, statistical analysis revealed a significant correlation between VP behavior and transition in the judgement of humanness (Fig. 3A) indicating that the latter depends on both cooperative behavior and how this behavior changes over time.

**Figure 3.**
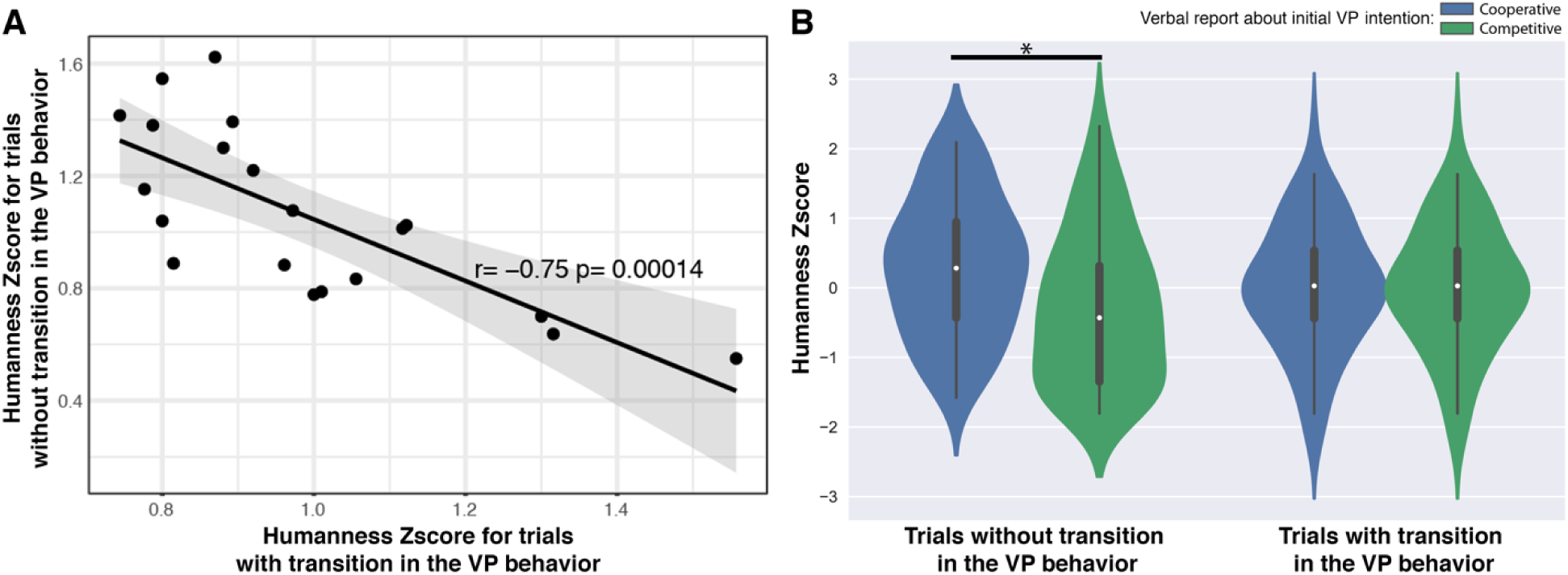
Humanness assessment during the interaction with the Virtual Partner. (**A**) Transitions in VP behavior modulated the subjective feeling of interaction with another human, measured by normalized Humanness score (Z-scores, raw values were between 0 and 1, respectively corresponding to judgement of VP as a Robot or a Human). Shaded area indicates 95% confidence interval. (**B)** Modulation of perceived humanness by the presence of transition in VP behavior and its initially perceived cooperativeness. Star indicates statistically significant differences (*p* < 0.05, corrected).

The 2×2×2 (VP Behavior [Cooperative/Competitive] by Transition [Yes/No] by Task [In-phase/Anti-phase]) repeated-measures ANOVA on humanness ratings showed a main effect of VP behavior (F(1, 19) = 7.022, *p* = 0.008, η^2^ = 0.050) as well as an interaction between VP behavior and Transition (F(1, 19) = 7.022, *p* = 0.008, η^2^ = 0.050) suggesting that a drop in humanness rating occurred when VPs were acting in a competitive fashion, and specifically when VPs sustained their competitive intention throughout the trial (see Fig. 3B).

### Right parietal activity during Human ∼ VP coordination

Power analyses on estimated cortical sources revealed well-known mu suppression in the upper alpha band over primary motor areas during movement (maximum difference over Precentral Left with t(19) = −5.58, *p* < 0.0001; Fig. 4A) and also revealed a joint suppression of high-alpha band (maximum difference over Parietal Superior Right: t(19) = −10.38, *p* < 0.001; Fig. 4B) and an increase in gamma band activity over right posterior cortex during active coordination with VP (maximum difference over Temporal Superior Right: t(19) = 2.95, *p* < 0.05; Fig. 4C). Interestingly, the emotional response of participants (as quantified by SPR) was also linked to a strong modulation of high-alpha activity in the same anatomical regions (maximum difference over Supra Marginal Right: t(19) = 3.55, *p* < 0.05; Fig. 4D).

**Figure 4.**
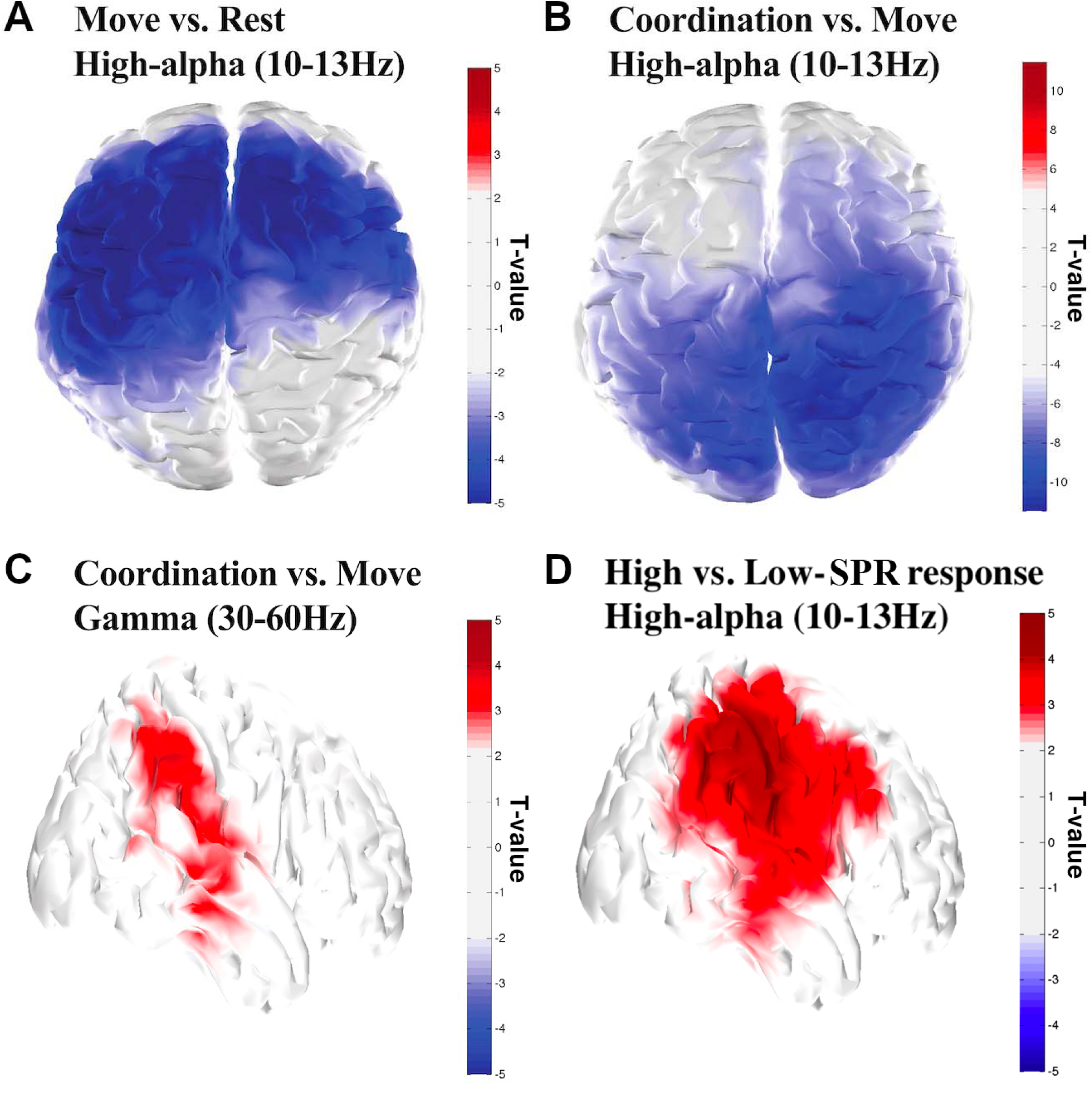
Modulation of spectral activity of cortical sources. High-alpha suppression over motor regions during execution of movement without VP (**A**), joint decrease of high-alpha activity (**B**) and increase of gamma activity (**C**) over right parietal cortex during interaction with VP, and modulation of high-alpha activity by the emotional responses of the participants as measured by Skin Potential Response (SPR) (**D**). Color indicates clusters of cortical sources which were significantly modulated in each contrast.

### Neural dynamics coordinated with self- and other behavior

Analysis of cortico-motor coherence with velocity of human movement (‘self’) revealed significant involvement of contralateral primary motor cortex (red colored region of the brain, maximum difference over Precentral Left: t(19) = 3.53, *p* < 0.005; Fig. 5). In contrast, when cortico-motor coherence was based on Virtual Partner (‘other’) velocity, an antero-posterior network was observed (blue colored regions of the brain, maximum difference over right cuneus: t (19) = 4.56, *p* < 0.0001; Fig. 5). Both brain networks, for self and other, span the fundamental frequency of movement and the first 2 or 3 harmonics (see distribution in the insert spectra, Fig. 5), roughly corresponding to the delta/theta bands.

**Figure 5.**
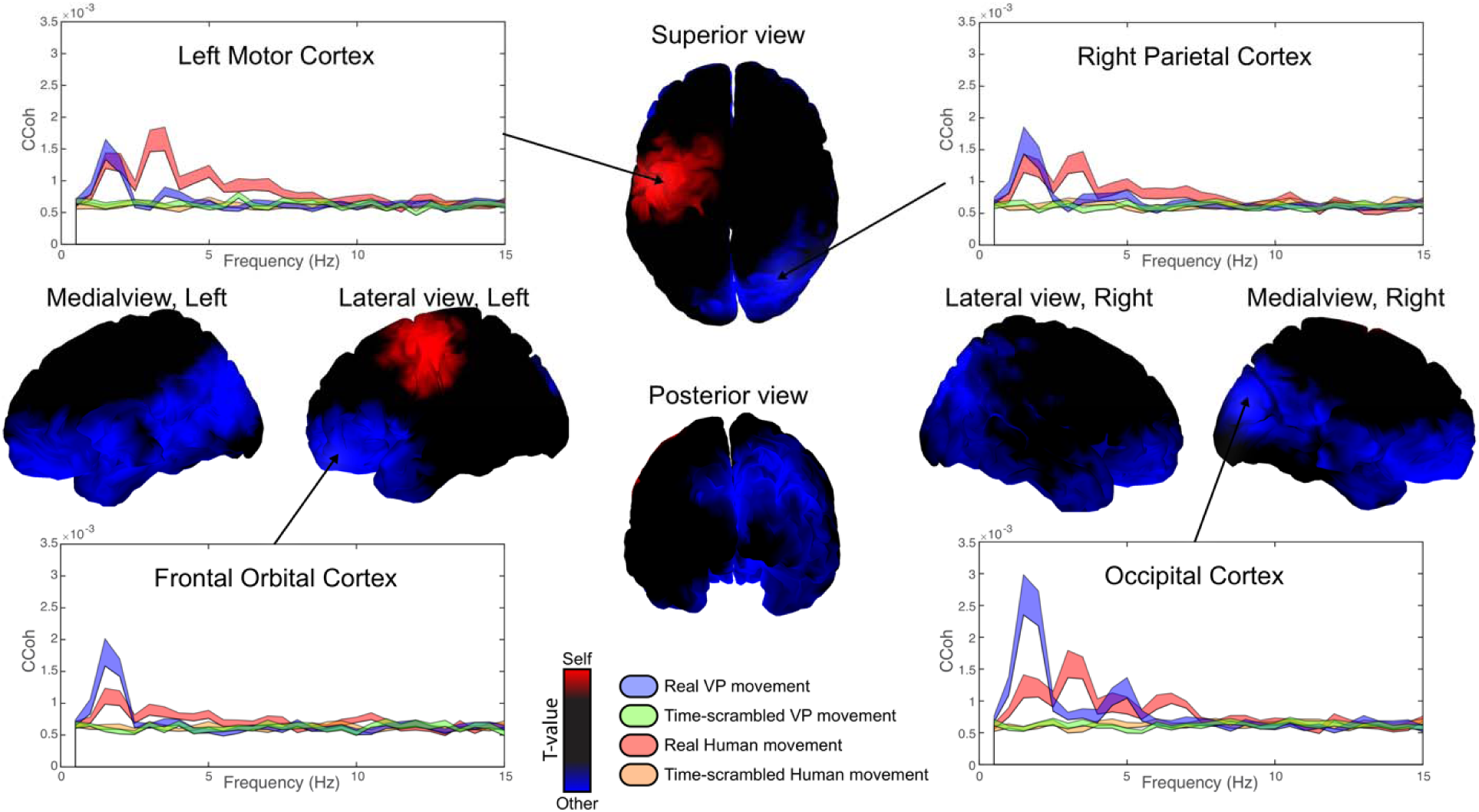
Self-other related cortico-motor coherence modulation. Graphs show cortico-motor coherence (Jerbi et al., 2007) across frequencies for key task-related brain structures. 3D brain figures represent statistical differences between cortico-motor coherence of human and VP movement velocities and neural activity in the delta/theta band (1-5Hz). Red and blue indicate sources which were statistically more associated with (respectively) self- or other-movement. Notice the increase for self-movement in the left motor cortex and for other-movement in right parieto-occipital, left frontal, and midline regions. Notice also the absence of coherence with time-scrambled movements (Green and Orange in the graphs). Shaded areas in 2D graphs represent standard errors.

Besides pointing to significant differences in corticomotor coherence between brain regions associated with self and other movement, we can investigate how the two networks are integrated via an analysis of their overlap (Fig. 6). Joint self and other movement coherence activities were found over right parietal areas (Purple) with four ROIs: Cuneus Right, Angular Right, Parietal Inferior Right, and Supra-Marginal Right (Fig. 6, pink color). Very few parts of the cortex were indifferent to both self and other’s movement (black color, restricted to superior frontal areas). The region that was specific for self-movement, again, was the contralateral motor cortex (red color). In contrast, a large expanse of the cortex was concerned with the other’s movement (blue color).

**Figure 6.**
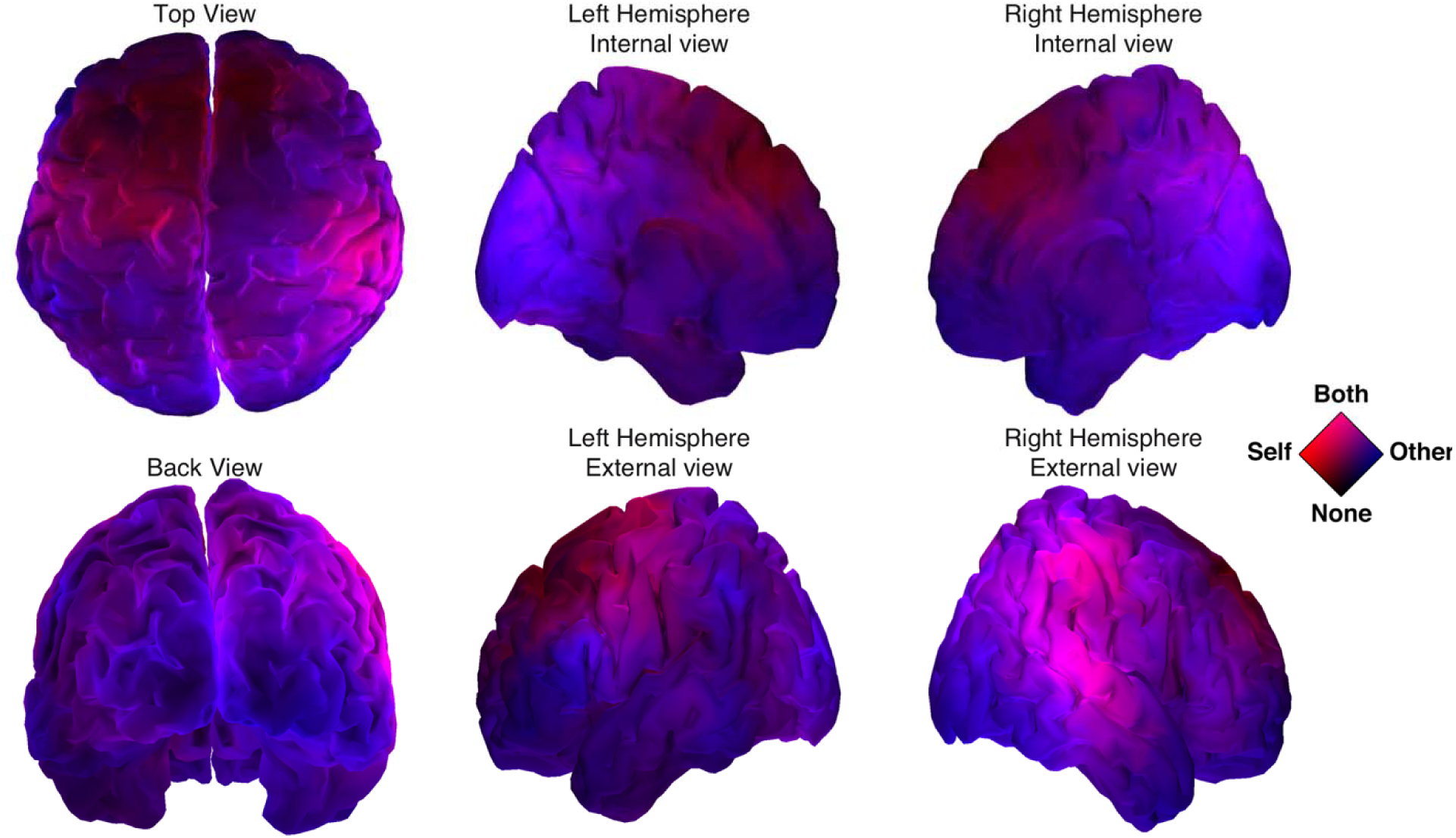
Overlap of self- and other-related brain networks. The networks related to self and other movement overlap on both right Temporo-Parietal Junction (rTPJ) and right Superior Temporal Sulcus (rSTS). Color stands for significance (-log(p)) of theta cortico-motor coherence contrasts (*p <* 0.05) for Real vs Scrambled behavior. Red and Blue color components code respectively for the “Self- vs. Scrambled” and “Other- vs. Scrambled” contrasts. Purple corresponds to cortical sources related to *both* Self and Other behaviors.

Modulation of functional connectivity by attribution of humanness and cooperation. Functional connectivity in the low-theta band was explored in order to understand how self- and other-related information may be related to cooperative/competitive behaviors or humanness judgement during social interaction. The two contrasts revealed a similar pattern: sensorimotor hubs in the posterior part of the brain, predominantly in the right hemisphere (e.g. Cuneus), were coordinated with anterior areas (e.g. Frontal Superior). Figure 7A-D illustrates the anatomical distribution of the most important changes due to coupling, revealing that both the attribution of humanness to the VP and cooperation of VP are associated with a coherence increase between posterior and anterior brain structures in the delta/theta band (1-5 Hz). Figures 7E-F provide further details on the brain areas involved in this increase (or decrease) of coherence according to the Tzourio-Mazoyer atlas.

**Figure 7.**
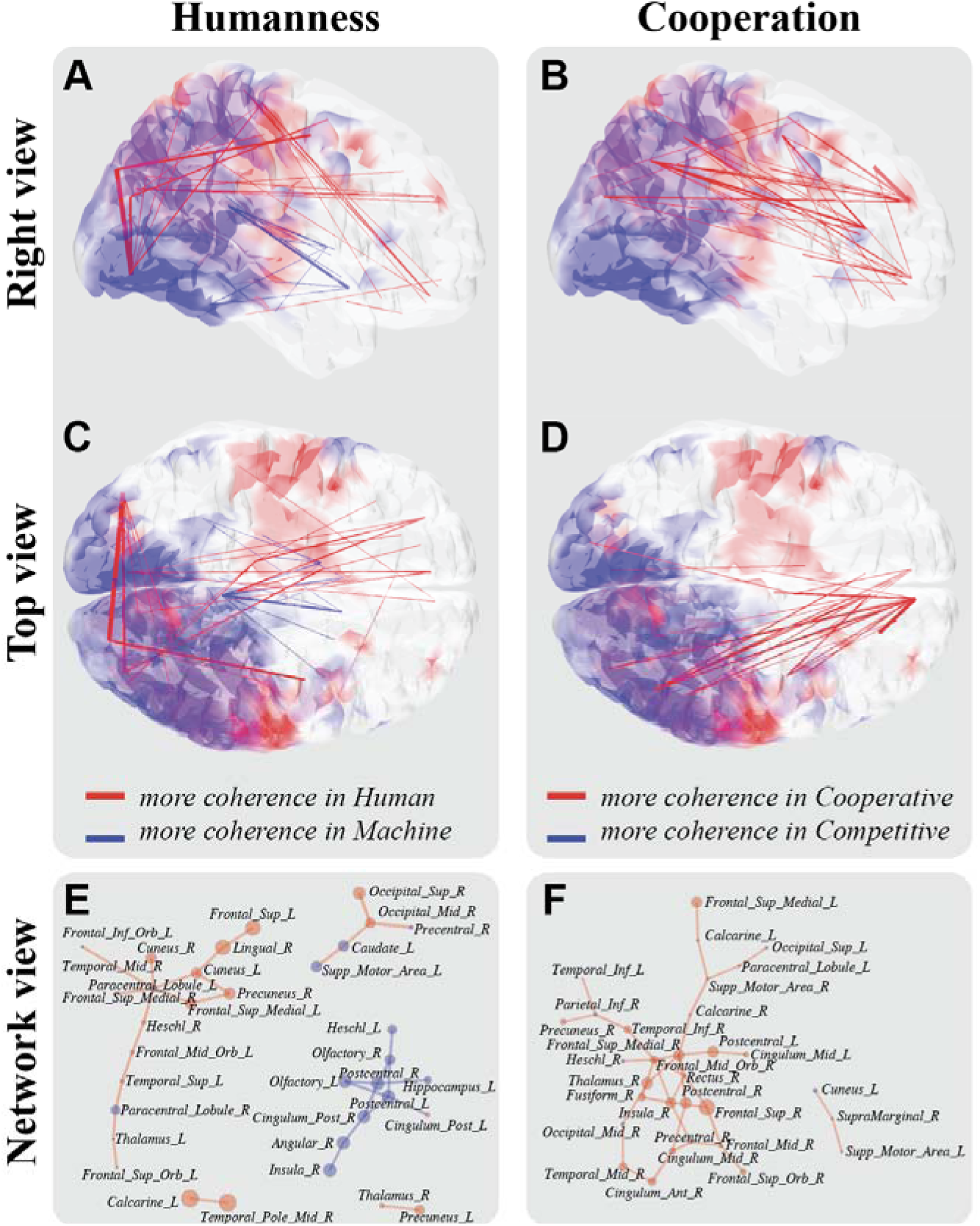
Right parietal cortex as a hub between self and other sensorimotor brain networks. Humanness attribution and cooperation are related to changes in large scale brain dynamics. Coherence between posterior and anterior brain structures in the delta/theta band (1-5 Hz) increased when participants judged the VP as human rather than machine (**A, C**, *p* < 0.05 non corrected) and when VP was cooperative rather than competitive (**B, D**, *p* < 0.05 non corrected). Panels (**E**) & (**F**) show respectively the networks of brain structures implicated in (**A**, **C**) and (**B**, **D**). Line width and circle size indicate the modulation strength of coherence and power in the theta band.

## Discussion

The present study, for the first time to our knowledge, used the Human Dynamic Clamp to investigate the neural underpinnings of social interaction and in particular how the brain integrates its own behavior with that of others to support attribution of humanness and intention. We recorded high-density EEG in humans interacting with a virtual partner parameterized to act in a cooperative or a competitive way. At the behavioral level, our results suggest a link between sensorimotor coupling and intention attribution as shown by the correlation between individual phase coordination scores (stability) and accuracy in detecting VP’s goals (i.e. to compete or to cooperate). At the brain level, we highlight the recruitment of networks associated with self-other integration, demonstrating a key overlap over right parietal areas.

### Attribution of intention to a Virtual Partner

At the behavioral level, our results first demonstrate how interaction with the Human Dynamic Clamp leads to successful attribution of intention. Indeed, participants accurately judged the intention of the VP despite the confounding factor of task difficulty arising from performance of in-phase vs anti-phase coordination (more difficult). The experimental protocol also led to attribution of humanness, though the partner was in all instances a computer program the fed input of which was from the human participant’s movement. Participants judged the VP to be human 47.3% of the time. Our data show that the presence of a mid-trial transition (see Figure 3) modulated attribution of humanness to the virtual partner and that global joint coordination (i.e. relative phase stability) was correlated with accurate attribution of intention to the VP. These behavioral results therefore confirm the ecological validity of coordinating with the HDC in real-time and demonstrate the influence of a virtual partner’s behavior on human social cognition. Previous human-machine interaction paradigms using gaze pattern have shown how humanness attribution can be modulated by contingency and task (Pfeiffer et al., 2011). Other psychosocial parameters such as gender similarity between a virtual partner and the human participant can impact subjective reports about behavior (Guadagno et al., 2007). Previous findings show how the HDC protocol allows a new exploration of relational patterns during real-time sensorimotor coordination and how the latter modulates subjective judgement (Kelso et al., 2009; Dumas et al., 2014a) and emotional responses (Zhang et al., 2016). Building upon those results and combining the HDC with high-density EEG recording, provides an opportunity to uncover the neural dynamics underlying a non-verbal Turing test during real-time social interaction.

### Coordination of self and other neural networks and the role of the right parietal cortex

At the brain level, source analysis reveals multiple brain networks involved in social coordination operating at different frequencies. Motor areas are recruited, as highlighted by a significant decrease of high-alpha/mu (10-13 Hz) power over contralateral and medial Rolandic regions during execution of movement compared to rest (Fig. 4A). This result is consistent with numerous reports of mu desynchronization during action execution (Salenius et al., 1997; Babiloni et al., 1999; Pfurtscheller et al., 2006; Dumas et al., 2014c; Hobson and Bishop, 2016). Furthermore, in line with previous studies targeting reciprocal exchange of movement information during social interactions (Dumas et al., 2010; Novembre et al., 2016), the present EEG results exhibit decreases in high-alpha power over the superior aspect of right posterior parietal areas during social coordination in comparison to similar movement produced without a partner (Fig 4B). The topography observed here is evocative of a body of work in monkey relating posterior parietal cortex to online control of visually-guided movement (Buneo and Andersen, 2006; McGuire and Sabes, 2011)--with the caveat that caution is warranted due to task (individual goal-directed actions vs. social coordination) and species differences (monkey vs. human). Nevertheless, the present results are suggestive that posterior parietal alpha suppression reflects the visual coupling governing movement coordination between partners, whether virtual or not. Our data also expands current knowledge on parietal dynamics during on-line social interaction (Tognoli et al., 2007; Era et al., 2018b) by highlighting an increase of source-resolved gamma power over right temporo-parietal areas during social coordination compared to solo actions (Figure 4C). Finally, interactions marked with high levels of emotional responses elicited increased alpha in a similar right parietal-temporal-insular complex (Fig. 4D; see Zhang et al., 2016). These results not only highlight the key role of right parietal areas in social coordination, but also point toward a link between sensorimotor neuromarkers and affective dimensions of human social cognition.

Previous research has identified involvement of right parietal cortex, specifically the temporo-parietal junction (TPJ) for the integration of self and other’s actions (Sowden and Catmur, 2015). TPJ is known to integrate inputs from subcortical (e.g. thalamus) and cortical (e.g. occipital, temporal, and prefrontal) regions (Decety and Lamm, 2007). Functional neuroimaging studies have repeatedly linked activity over right TPJ (rTPJ) with socio-cognitive processes (Decety and Chaminade, 2003), including Theory of Mind and empathy (Jackson et al., 2006), and joint-attention (Redcay et al., 2010). Moreover, research by Bzdok and colleagues (2013) has revealed the presence of two distinct clusters within rTPJ that are associated with either attentional processes or social cognition. Interfering with brain activity using Transcranial Magnetic Stimulation has shown a causal role of rTPJ in numerous processes relevant to social interaction, ranging from self-centered (i.e. body ownership; Tsakiris et al., 2008) and other-centered socio-cognitive processes (mentalizing and Theory of Mind; Bardi et al., 2017) to self-other integration in imitative actions (Sowden and Catmur, 2015). Crucially, enhancement of rTPJ activity (using tDCS) improves online interactions by boosting the ability to switch between self and other representations in both perspective-taking and the control of imitation (Santiesteban et al., 2012). Furthermore, Era et al. (in review) suggest that rTPJ supports the ability to perform joint movements only when self and other movement planning overlaps. The present research adds the understanding that rTPJ is the site of integration of corticomotor frequencies in the theta range for self and other, and that it is accompanied by higher frequencies in the gamma band for sensorimotor coordination.

Cortico-motor coherence in the theta band also reveals how right parietal sources are coordinated with shared movement velocity (Figure 6). The similar topography (consistent with the location of rTPJ) of the gamma increase, the alpha decrease and the cortico-motor coherence in the theta band suggest a multifaceted integration of self and other behaviors and a polyvalent role of right parietal areas in supporting sensorimotor coordination. Alpha desynchronization over parietal sites has been linked previously to ‘gating’ information (Varela et al., 1981; Busch and VanRullen, 2010). Activity in the theta band is usually associated with long-range connectivity and large-scale coordination of distant cortical areas (Moreau et al., 2018), a finding that led Zhang and colleagues (2018) to describe theta waves as ‘traveling’ waves, crucial for working memory tasks. Conversely, activity in the gamma band is often associated with the local processing of information (Fries, 2009). Much previous research has reported a coupling of theta and gamma activity over similar areas thought to form a neural code for representing sequential order among numerous elements (Canolty and Knight, 2010; Lisman and Jensen, 2013). Here, the co-localization of the theta hub between self- and other-network and the gamma activity associated with active social coordination points toward a functional cross-frequency link uniting the two processes.

Social interactions are not solely orchestrated by parietal regions. Moreau et al. (2018) showed an increase of occipito-temporal activity when a virtual partner performed unexpected actions during motor interactions, highlighting a role of occipito-temporal areas in integrating the behavior of others. In accordance with this, we find that an increase of cortico-motor coherence over posterior sites is related to the partner’s movement (see Figure 5). The present connectivity analysis revealed a significant increase of cortico-cortical coherence between bilateral occipital areas when participants declared they were interacting with a human partner (Fig 7C and 7E).

### Brain dynamics of social embodiment

Understanding the intentions of others is a crucial feature of effective social interaction. The present behavioral results highlight a correlation between sensorimotor performance (i.e. behavioral coupling with the VP) and correct attribution of the VP’s intentions. Corresponding brain analysis reveals how anterior areas coordinate their activity with frontal and prefrontal areas, the latter generally recruited in decision-making tasks regardless of the presence of others (Tomlin et al., 2006; Campbell-Meiklejohn et al., 2017; Shaw et al., 2018; Thornton et al., 2019). Our results thus appear to fit the nexus model of rTPJ (Carter and Huettel, 2013), which proposes that initial processing of social information by occipital and occipito-temporal areas precedes the integration/segregation of relevant information in rTPJ.

Cooperation is central to human societies and has played a key evolutionary role in human social behavior (Barrett et al., 2010; Dunbar, 2011; González-Forero and Gardner, 2018). Our results lend support to the idea that there is an imbalance between self and other neural dynamics, the other coming before the self (Graziano and Kastner, 2011). Taken together, the present findings call for going beyond the social brain *per se* to embrace a more integrative perspective where sensorimotor abilities and accurate intention attribution are part and parcel of the same self-organizing coordination dynamics that grounds social awareness and cognition (Oullier and Kelso, 2009; Kelso et al., 2013; Dumas et al., 2014b).

## Acknowledgements

This work was supported by the National Institute of Mental Health grant MH080838, the National Science Foundation grant BCS0826897, and The Davimos Family Endowment for Excellence in Science. G.D. also acknowledges support from the Innovative Medicines Initiative (IMI) during completion of this work. We thank the Brainstorm Team for their software (http://neuroimage.usc.edu/brainstorm/) and G. de Guzman for help in implementing the Human Dynamic Clamp. We also thank M. Zhang for her support in the earlier stages of this research.

